# Whole-brain dynamics of articulatory, acoustic and semantic speech representations

**DOI:** 10.1101/2024.08.15.608082

**Authors:** Maxime Verwoert, Joaquín Amigó-Vega, Yingming Gao, Maarten C. Ottenhoff, Pieter L. Kubben, Christian Herff

## Abstract

Speech production is a complex process that traverses several representations, from the meaning of spoken words (semantic), through the movement of articulatory muscles (articulatory) and, finally, to the produced audio waveform (acoustic). In our study, we aimed to identify how these different representations of speech are spatially and temporally distributed throughout the depth of the brain. By considering multiple representations from the same exact data, we can limit potential con-founders to better understand the different aspects of speech production and acquire crucial complementary information for speech brain-computer interfaces (BCIs). Intracranial speech production data was collected of 15 participants, recorded from 1647 electrode contacts, while they overtly spoke 100 unique words. The electrodes were distributed across the entire brain, including sulci and subcortical areas. We found a bilateral spatial distribution for all three representations, although there was a stronger tuning in the left hemisphere with a more widespread and temporally dynamic distribution than in the right hemisphere. The articulatory and acoustic representations share a similar spatial distribution surrounding the Sylvian fissure, while the semantic representation appears to be widely distributed across the brain in a mostly distinct network. These results highlight the distributed nature of the speech production process and the potential of non-motor representations for speech BCIs.

## Introduction

Neuroimaging studies have shed light on various components of language representation in the brain for many years. Recently, the representation of semantic information has been more comprehensively mapped thanks to the aid of large language models^1–3^. Conversely, the definitions of Broca’s area, traditionally associated with speech production, and Wernicke’s area, traditionally linked to language comprehension, have been up for debate due to emerging evidence that challenges their classical boundaries and functions^4–9^. These findings, along with results from more data-driven analyses and neural data with high temporal resolution from deeper brain areas, suggest that the canonical language map^10,11^ may be changing.

Different components of speech have been mapped to specific, but also many shared regions in the brain. The left inferior frontal gyrus (IFG), left precentral gyrus, left superior temporal gyrus (STG), left fusiform gyrus, right STG, and bilateral supplementary motor area (SMA) have been identified as “core” cortical language regions^10^. Speech production studies have shown strong left-lateralization, while speech perception appears to be more bilateral^1,10^. More specifically, the ventral “comprehension” stream, involving auditory and temporal cortices, is thought to be organized bilaterally, while the dorsal “articulatory” stream, involving mostly frontal and motor cortices, is thought to be strongly left-hemisphere dominant^11^. However, more recent evidence suggests that even the articulatory representation of speech production occurs bilaterally^12,13^, as well as other speech components^4^. Moreover, speech studies have largely overlooked potentially important deeper cortical and subcortical areas such as the insula^14^, basal ganglia^15,16^, thalamus^17^ and hippocampus^18^.

Traditionally, studies investigating speech, or language in general, have employed meticulously designed hypothesis-driven approaches to focus on isolated aspects of language, often in speech perception. Using advances in Natural Language Processing and Deep Neural Networks, data-driven approaches now allow us to explore characteristics of speech in natural continuous speech and language^1,2,19^. Another recent departure from traditional approaches is a shift from encoding to decoding^20^ studies of speech. Being able to decode speech from neural data alone can ultimately be used for speech neuroprostheses, also termed speech brain-computer interfaces (BCIs). Speech BCIs are currently developed with the ultimate goal of restoring communication in individuals who have lost their ability to speak^8,21–26^.

One of the approaches to decoding speech is through articulatory features. These features represent the movements of articulatory organs (such as the larynx, tongue, and lips). Previous studies have mapped the neural representation of speech articulators to areas such as the ventral sensorimotor cortex^27–29^ and the dorsal precentral gyrus^30^. Another approach is to extract spectrotemporal features of the recorded speech audio to obtain an acoustic representation of speech. These features have been used to decode and synthesize speech from the inferior frontal and motor cortex^31^, the STG^32^ and from a combination of multiple distributed regions^33,34^. The articulatory and acoustic representations have both, separately, already led to successful BCI communication in an individual with paralysis due to a brain-stem stroke using a device on the sensorimotor cortex^35^. Combining these two approaches has the potential to further improve BCI performance^36,37^. While these studies have decoded speech from motor or acoustic representations of speech, individuals with diseases that cause impairments in the creation of motor representations, such as with aphasia, or in the generation of audible speech, such as with locked-in syndrome, might benefit from the decoding of more semantic representations of speech. The semantic representation captures the underlying meaning of words or sentences. Through the aid of large language models, it has been shown that we are able to capture such semantic representations as well, using both functional magnetic resonance imaging^1,3^ (fMRI) and intracranial electrophysiology^2,19,38^. These studies found semantic representations spread widely across the brain. The question remains whether these different speech representations share a similar or distinct network of brain regions.

While exploring these different speech representations is crucial, understanding the strengths and limitations of the methods used to study them is equally important. fMRI is a useful method to study language representations as it is non-invasive and can capture whole brain dynamics, however, it suffers from various sources of noise (e.g., head motion, loud scanner noise) and is particular sensitive to the motion artifacts induced by speaking. Furthermore, fMRI is lacking in temporal resolution, which is crucial for certain aspects of speech. Intracranial electrophysiology, on the other hand, is characterized by its high temporal resolution, although the spatial resolution depends on the specific electrode device and implantation planning. Stereo-electroencephalography (sEEG), specifically, is routinely used for functional mapping and to determine the seizure onset zone in epileptic patients^39^. The sEEG electrodes are shafts with individual contacts that reach subcortical areas and are typically implanted in many different locations at once. In the current study, we analyze sEEG data from multiple participants with distinct and overlapping electrode locations which allows for a high overall spatial resolution, a brain-wide coverage, together with high temporal resolution. This is ideally suited to study the widespread process of speech production.

Language thus appears to be encoded in a wide variety of brain regions, although the methods used to study it also vary in terms of speech features, analyses and technology. We aimed to provide an overview of how multiple speech representations are neurally distributed throughout the brain, using similar analyses on the same high-resolution data. We focus on the three representations of the speech production process that are most commonly used for the purpose of speech BCIs^26^, namely articulatory, acoustic and semantic representations. We utilize data-driven approaches to extract and compare these multiple speech features from the same exact data in a continuous manner. We employ an Acoustic-to-Articulatory Inversion Recurrent Neural Network^40^ to estimate the movements and positions of articulatory organs during speech production to generate an articulatory representation, a Fourier transform to capture auditory properties of spoken speech as an acoustic representation and we utilize word embeddings from a Word2Vec auto-encoder^41^ for a semantic representation (Fig. 1). The results shed further light on the overall speech production network, including deeper regions, and are discussed in the context of speech neuroprostheses.

**Figure 1.**
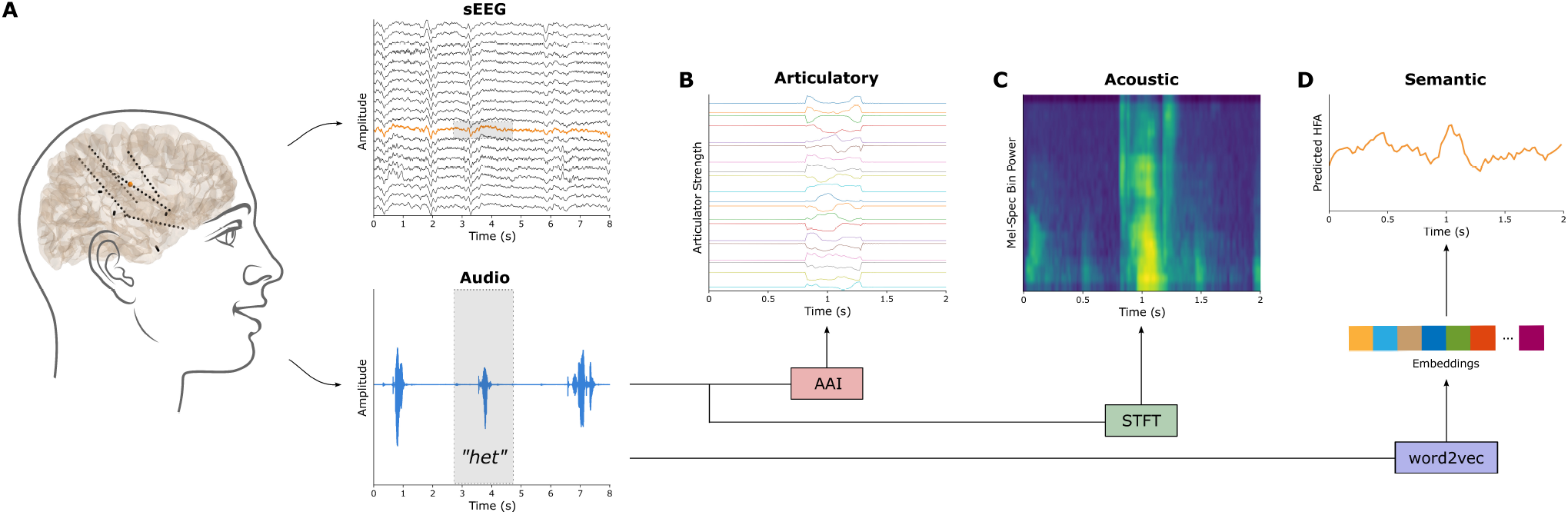
Extraction of the three speech representations. **A)** The dataset contains synchronized sEEG and audio data during speech production. Highlighted is one single channel’s neural signal (also in orange) and the audio signal of one word. **B)** The articulatory trajectories (N=20) were estimated using an acoustic-to-articulatory inversion (AAI) LSTM neural network. **C)** The acoustic representation in the form of mel-spectrograms was calculated using short-time Fourier transform (STFT) with N=23 filterbanks. **D)** The semantic embeddings (160 dimensional) were extracted for each word using a word2vec model and a linear regression was subsequently trained to predict the neural timeseries.

## Results

### Differential distribution of speech representations

For all three speech representations (articulatory, acoustic and semantic), there are significant channels present in both hemispheres (Fig. 2A). Whereas the total number of significant channels is similar between the three representations (Fig. 2B), there are relative differences between the two hemispheres (Fig. 2C). Since there were a greater number of channels implanted in the left hemisphere (937) than in the right hemisphere (710), we looked at the percentage of significant channels within the hemispheres. Percentage-wise, there are more significant channels in the left hemisphere for the articulatory and acoustic representations. While the articulatory and acoustic representations are largely overlapping in their spatial distribution (74.22% out of all articulatory and/or acoustic channels), there appears to be a relatively stronger tuning for articulatory features in the left hemisphere versus a relatively stronger tuning for acoustics in the right hemisphere (Fig. 2C). The articulatory-acoustic overlap can be seen in yellow in Figure 2A and Figure 2B. The semantic representation, on the other hand, has little overlap with the other two representations (16.52% overlap out of all semantic channels).

**Figure 2.**
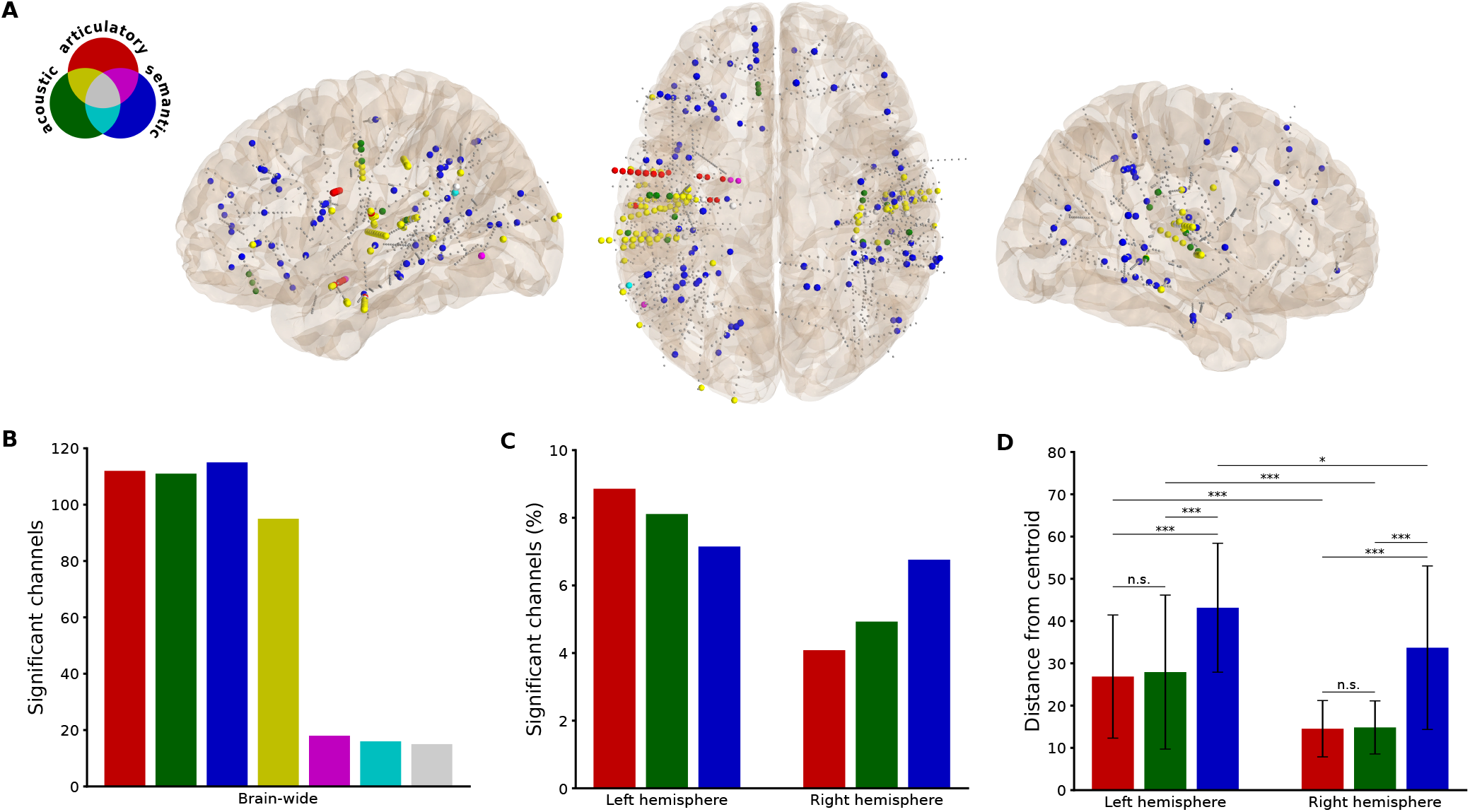
Differential distribution of speech representations. **A)** Significant channel locations with colors representing each representation and their overlap, smaller gray dots represent non-significant channels. **B)** Total number of significant channels for each representation and the overlap between representations. **C)** Percentage of significant channels for each representation, separated by hemisphere. **D)** The mean Euclidean distance between each channel and a centroid for each representation, calculated per hemisphere. Significance was tested using independent samples t-tests (*** p < 0.001, * p < 0.05, Bonferroni-corrected).

For each representation, there was a significant difference in the average distance of significant channels from their centroid between the two hemispheres (Fig. 2D), indicating that all representations are more widely distributed across the left than across the right hemisphere. This difference could potentially be due to the sampling, since the electrode coverage is not exactly mirrored between the hemispheres (along with more contacts in the left hemisphere overall). Therefore, we additionally calculated the distance between all recorded channels and their centroid in each hemisphere. This revealed no difference between the left (38.47 ± 16.69) and the right (37.25 ± 17.57) hemisphere overall (*t*(1645) = 1.44, *p* = 0.15). The difference is therefore not likely due to the electrode coverage alone. The semantic representation is also more widely distributed across the brain than the other two representations, in both hemispheres (Fig. 2D). Importantly, this holds true within individuals (14/15 participants). Only for one individual the articulatory-selective channels were spread farther than the semantic ones (Fig. S2). Together, these findings indicate distinct and widespread processing of the meaning of words.

### Anatomical contributions to speech representations

Since the overall distribution covers many different anatomic areas, including contacts located in white matter tracts (80 out of the 224 (35.7%) overall significant channels), we next looked at how specific regions-of-interest (ROIs) may contribute differently to the speech representations (Fig. 3). We find the strongest contributions from auditory and motor regions, with a preference for acoustic features in the auditory cortex and a slight preference for articulatory features in the motor cortex. Interestingly, these regions were strong contributors in both hemispheres. However, the semantic representation appears to be stronger in the right hemisphere in these primary regions specifically.

**Figure 3.**
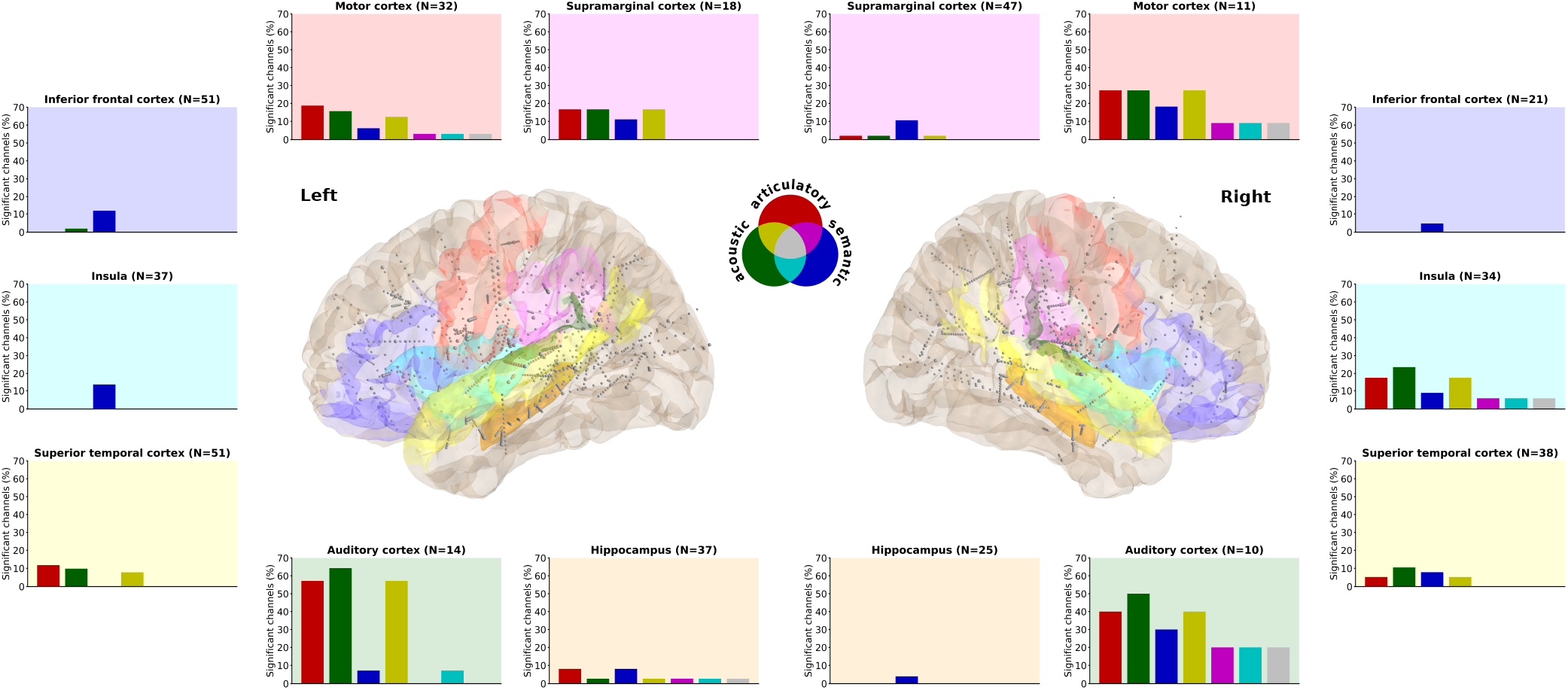
Anatomical contributions to speech representations. Seven language-related anatomical areas within each hemisphere are highlighted. The percentage of significant channels (out of the total number of channels in that area) for each representation and their overlap is displayed in the corresponding bar plot. Background colors in the bar plots match and indicate the regions in the brain plots where all channels are depicted in grey, the significant ones are slightly larger in diameter.

The semantic representations are present in nearly all the ROIs, although to a relatively small degree. It is the only representation we find in the inferior frontal gyrus, the left insula and the right hippocampus. The left hippocampus also has a small articulatory and even smaller acoustic representation. The right insula, on the other hand, has stronger articulatory and acoustic representations. There are weak representations in the superior temporal cortices, relative to the auditory regions. Finally, the supramarginal cortex appears to be involved specifically in the left hemisphere.

### Temporal dynamics of speech representations

To explore the brain-wide temporal dynamics, including channels that are not represented in ROIs, we grouped the results into three time segments. They represent time windows prior to speech alignment (−200 until -50 ms), surrounding speech alignment (−50 until +50 ms) and after speech alignment (+50 until +200 ms) for the articulatory and acoustic representations. For the semantic representation, the two-second trial time points were equally divided, however, since speech onset latency is roughly between 500-600 ms^42^, the middle segment is similarly expected to involve most of the word production. Spatially, we find a relatively stable pattern in the right hemisphere for the articulatory-acoustic representations and there appears to be a posterior to anterior pattern in the left hemisphere for the semantic representation (Fig. 4A). Notably, there is one articulatory-selective electrode shaft in the left precentral or motor cortex specifically prior to and after speech onset and one acoustic-selective electrode shaft in the left postcentral or sensory cortex specifically after speech onset.

**Figure 4.**
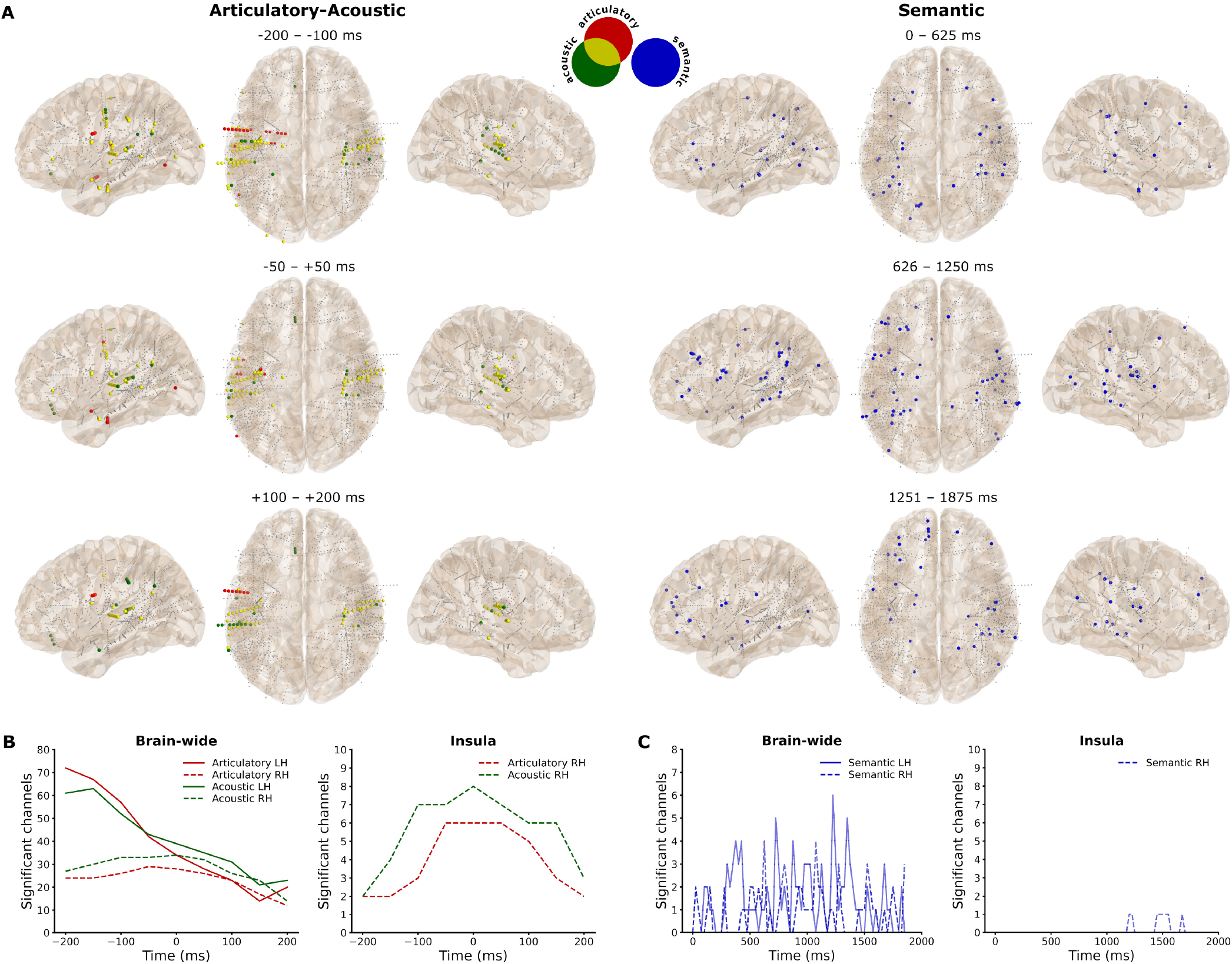
Temporal dynamics of speech representations. **A)** Significant channel locations with colors representing the representation and the articulatory-acoustic overlap, divided into time segments. Smaller gray dots represent non-significant channels. **B)** Number of significant articulatory- and acoustic-selective channels across time-frames from all (brain-wide) contacts and from the right insula. **C)** Number of significant semantic-selective channels across time-points from all (brain-wide) contacts and from the right insula.

The amount of significant channels dynamically changes across timeframes, especially in the left hemisphere (Fig. 4B). In the left hemisphere, most significant channels were found 200 ms prior to speech onset, after which the number quickly declines to a level similar to that of the right hemisphere at around +50ms. In contrast, in the right hemisphere, the number of significant channels is relatively stable until it drops after +50 ms. Notably, in the left hemisphere, the peak for the acoustic representation is one timeframe later (−150 ms) than for the articulatory representation (−200 ms) and there is a switch around 50 ms prior to speech onset where the acoustic representation becomes more prominent than the articulatory representation. We find similar patterns in most of the ROIs (Fig. S3), although the right hemisphere insula stands out. Here we find a steep increase in the articulatory- and acoustic-selective channels until around speech onset, which drops more slowly afterwards for the acoustic representation (Fig. 4B).

The largest number of significant channels is roughly around the middle of the speech trial for the semantic representation (Fig. 4C). The channels fluctuate across time and locations and between hemispheres. The spatial results, as mentioned before, indicate that the locations fluctuate from more posterior to more anterior in the left hemisphere, although this is not apparent in the right hemisphere (Fig. 4A). Finally, for appropriate comparison we also highlight the right insula for the semantic representation, which is also involved relatively late during or after speech production (Fig. 4C).

## Discussion

In this work, we explored the brain-wide spatial distribution and temporal dynamics of articulatory, acoustic and semantic speech representations. The results indicate more widespread and temporally dynamic contributions from the left compared to the right hemisphere, in all representations. This is in line with the broad literature on the left-hemisphere dominance in speech production^10,11^. However, we do note strong contributions from the right hemisphere as well, especially from the primary auditory and motor regions as well as from the right insula. Thus, the results also align with the notion of bilateral involvement in speech production^12,13^. The temporal dynamic results indicate that the majority of articulatory- and/or acoustic-selective channels are active well before speech onset. This finding is especially pronounced in the left hemisphere, even in auditory regions. This could reflect the communication between different areas for speech preparation. Interestingly, the neural patterns were temporally stable in the right hemisphere. This activity could potentially reflect feedback control, for which the right hemisphere has previously been implicated^43^.

The articulatory and acoustic representations share a similar neural distribution, most prominently within and surrounding the Sylvian fissure. Similar overlap has been found in earlier studies using fMRI^44^, sEEG^13^ and both representations were decodable with a chronic implant on the somatosensory cortex^35^. The semantic representation has a widespread neural distribution, which is mostly distinct from the articulatory-acoustic network, which is also consistent with a previous fMRI study^44^ on speech comprehension. This indicates that semantics may indeed operate in distinct networks. However, it is worth noting that there was more overlap between the representations in the right hemisphere, especially in the auditory cortex.

The semantic representation encompassed temporal, parietal and frontal regions. This distribution is in line with previous studies investigating semantic processing in speech comprehension^1–3,44^, indicating that the same semantic network is engaged in both speech comprehension and production. A key difference lies in the semantic context, while earlier studies involved listening to sentences or stories, our work focused solely on speaking isolated words without a broader semantic context. We therefore show that it is possible to uncover semantic processing in speech production, even with limited semantic context, a small amount of data, and without an explicit semantic task. The distributed results align with the ‘spokes’ part of the hub-and-spoke theory of semantic cognition^45^. They position the mediating ‘hub’ at the bilateral anterior temporal lobe, which unfortunately was not sampled sufficiently in our recordings to comment on further. In contrast to the articulatory-acoustic representations, there was a similar relative number of semantic-selective channels between the hemispheres, although we noted a more widespread and dynamic temporal pattern in the left hemisphere, moving from early posterior to later anterior regions. This may be explained by previous findings that the left hemisphere primarily represents language-mediated semantic knowledge, whereas the right hemisphere encodes more perceptually based sensory-motor conceptual representations^46^. This might also explain the relatively larger contribution in the motor and auditory regions in the right compared to the left hemisphere. Future research may explore whether the temporal pattern could additionally vary depending on the input domain (visual vs. auditory) and its corresponding processing hierarchy.

The inferior frontal cortex, also termed Broca’s area, was almost exclusively and weakly (to a similar degree as other regions-of-interest) correlated with the semantic representation. This region was previously found to be specifically active for overt word generation and not for simple (meaningless) overt sound reading^47^ and may be more involved in the higher-order executive control of semantic knowledge than its representation per se^45^. More involvement might be observed when more semantic context needs to be processed, i.e., in word or sentence generation. The lack of clear articulatory and acoustic representations in this region further shows its indirect connection to the low-level articulatory and acoustic features of speech, limiting its use for an articulatory-based speech neuroprosthesis^8^.

The superior temporal cortex showed low selectivity for any speech representation, in contrast to very high selectivity in the auditory regions. While the superior temporal gyrus (STG) is known to play a key role in speech comprehension^48^, its role in speech production is less clear. The posterior portions of the STG and superior temporal sulcus (STS) have been more specifically implicated in speech production^43^. However, this portion may be masked by the more anterior regions of our relatively large region-of-interest (ROI) and was partially (the ‘planum temporale’) included in the auditory cortex ROI. On the other hand, the STG has been found to entail intermediate acoustic-to-semantic sound representations that neither acoustic nor semantic models could account for^49^. This may similarly be the case for the opposite direction (semantic-to-acoustic representations), which could explain the relatively weak correlations with our representations. Another study found a special role of the right posterior STS (which may be closer to our auditory cortex ROI) in matching auditory expectations with spectrotemporal processing from auditory feedback during speech production and propose its involvement in the internal representation of speech^50^. For a speech neuroprosthesis, the question remains whether the auditory cortex remains involved when speech is only imagined instead of overtly produced. The temporal dynamics results indicate that even these regions, especially in the left hemisphere, are active well before speech onset. Other studies have found speech-related information in auditory regions even with imagined speech^50–52^. Together, these results suggest that the activity may indeed persist even when no sound is produced, which is important for the development of speech neuroprostheses for individuals who are unable to produce sounds.

Interestingly, the right insula showed relatively strong involvement in the speech production process, particularly for the acoustic and articulatory representations. While this area seems an interesting candidate for a speech neuroprosthesis^53^, the temporal dynamics, revealing most involvement around speech onset rather than before, suggest that it may not be a good predictor of speech. The insula is an integration hub linking many different cortical and subcortical regions and its activity may reflect any of a large number of functions^54^. However, since distinct regions of the insula have been shown to play different roles in speech and language processing^14,55^, its potential may still need to be explored in further detail.

The hippocampus has previously been shown to be involved in language processing^18^, more specifically, in navigating semantic representations^56^. However, we do not see a large contribution from the hippocampi in our results. This may, in part, be due to the fact that we did not explicitly probe semantic relations or memory, as only isolated words were produced as opposed to sentences or longer narratives. Additionally, our participants may have abnormal hippocampal activity due to their epilepsy^57^. Another recent sEEG study did find a large contribution from the hippocampus in decoding the production of vowels^58^, however, this was in combination with micro-wires. They did not find good decoding on the sEEG high-frequency local field potential on its own, which is similar to our work. Alternatively, previous findings linking the hippocampus with language were found in low-frequency activity^59^, which we did not include in our current work.

The supramarginal cortex was particularly involved in left hemisphere. Similarly, stimulation of this region was recently found to cause speech arrest with a short latency^60^. Both words and pseudowords were also recently found to be decodable with both overt and covert (imagined) speech using chronic implants on the left supramarginal gyrus^25^. This region is thought to potentially encode phonetics, however, they also found it to be activated during a visual imagination strategy and, separately, for decoding hand gestures. We also found all three speech representations in this region, together suggesting that it may serve as a potential area for a flexible and multi-purpose neuroprosthesis. However, they also noted a large difference between their two subjects in terms of decoding and anatomical folding of the region, suggesting the importance of individually localizing the specific area related to speech. While we did not find much representation other than semantics in the right hemisphere, it has to be noted that the specific electrode locations within the region, as well as in any other ROI, may differ between the hemispheres. Finally, even though exploring the white matter goes beyond the scope of the current paper, a large percentage (35.7 %) of significant channels were located in white matter. This suggests that we may be able to tap into these deeper communication highways directly.

## Conclusion

The articulatory and acoustic speech representations share a similar bilateral neural distribution surrounding the Sylvian fissure, most prominently in the left deeper auditory regions prior to speech onset. The semantic representation is spatially distinct from the others in a widespread frontal-temporal-parietal network, similar to what is seen in speech comprehension. While the primary auditory and motor regions show bilateral involvement, the insula was particularly correlated in the right hemisphere and the supramarginal cortex particularly in the left hemisphere. In combination, our results shed further light on the distributed and widespread networks involved in the speech production process and open the path for speech neuroprostheses based on differential representations of speech.

## Methods

### Participants

This study includes 15 Dutch-speaking participants (6 female, 9 male, age 36.9 ± 13.7 (range 16-60) years) who were implanted with sEEG electrodes to investigate the onset zone of their epilepsy. They agreed to participate on their own volition and provided written informed consent. Electrodes were implanted purely based on clinical needs. The Institutional Review Boards of both Maastricht University (METC 2018-0451) and Epilepsy Center Kempenhaeghe approved the study.

### Task

The task consisted of 100 unique Dutch words, individually presented on a screen for 2 seconds. During this time, the participant read and spoke the word out loud once. This was followed by a 1-second inter-trial interval with a fixation cross. Of note, the data of sub-10 only contains 95 words due to a technical issue.

### Data acquisition

The neural recordings were obtained using either one or two Micromed SD LTM amplifiers (Micromed S.p.A., Treviso, Italy), each equipped with 64 channels and operating at a sampling rate of 1024 Hz or 2048 Hz. Subsequently, the signal sampled at 2048 Hz was downsampled to 1024 Hz. The sEEG electrode shafts (Microdeep intracerebral electrodes; Dixi Medical, Beçanson, France) featured a diameter of 0.8 mm, a contact length of 2 mm, and an inter-contact distance of 1.5 mm. The number of contacts on a given electrode shaft varied, ranging from 5 to 18, with an overall count of implanted shafts ranging between 5 and 19 per participant. The data was recorded using ground and reference electrodes typically located in white matter regions that did not show epileptic activity. For audio data acquisition, the onboard microphone of the recording laptop (HP Probook) was utilized at a sampling rate of 48 kHz. LabStreamingLayer^61^ ensured synchronization among neural, audio, and stimulus data (Fig. 1A).

### Electrode localization

A custom version of the img_pipe^62^ Python package was used to localize electrodes alongside Freesurfer. The processes involved the co-registration between a pre-implantation T1-weighted anatomical magnetic resonance imaging (MRI) scan and a post-implantation computerized tomography (CT) scan, manual identification and inspection of contacts, and a non-linear warping to an average brain (MNI152) which was used only for visualisation purposes and the distribution metric. Anatomical labels were extracted using the Fischl atlas^63^ for subcortical areas and the Destrieux atlas^64^ for cortical areas. There were 1647 recorded electrode contacts (937 left hemisphere, 710 right hemisphere), besides the reference and ground contacts, spread across 128 unique anatomical labels (Fig. S1).

### Preprocessing

The data was re-referenced to an electrode shaft reference, in which the average of all other electrodes on a shaft is subtracted from each electrode on that shaft. In each channel, the neural signal underwent filtering to isolate broadband high-frequency activity (70-170 Hz) using an IIR bandpass filter with a filter order of 4. Subsequently, the Hilbert envelope was extracted for further analysis. To mitigate interference from the 50 Hz line noise, two IIR bandstop filters with a filter order of 4 were implemented. These filters were applied in both the forward and backward directions to eliminate any potential phase-shifts. Since neural activity precedes (in case of production) or succeeds (in case of perception) behaviour, we shifted the neural and audio data ranging between -200 and +200 ms in steps of 50 ms. These time-shifted signals were added as ‘timeframe’ features for the articulatory and acoustic data analyses.

### Signal analysis

#### Articulatory

The neural signal was averaged across 50 ms windows with a 10 ms frameshift. To generate the articulatory trajectories from the audio data, we first processed the audio by manually filtering out background noises unrelated to speech. We then modified the experimental markers to ensure that the markers indicating the start and end of the word trials always included the entire audio of the trial’s word. The modification was necessary as participants sometimes initiated speech late and did not finish within the allotted trial time. Three words in total were not spoken at all and therefore discarded from further analysis in the pipeline (Table S1). The markers were further adjusted to ensure a crucial 100 ms of silence at both the beginning and end of the utterances. This step is essential as the quality of the reconstructed articulatory trajectories is susceptible to the acoustic properties of utterances, especially the initial and final segments, particularly for plosive sounds.

The acoustic-to-articulatory inversion (AAI) pipeline contains different machine learning models trained using 16 kHz audio segments. Thus, we decimated our signal from 48 kHz to 16 kHz. We then cut the audio at the redefined markers and created segments for each word. The AAI pipeline also required an estimated syllable count for each word, which we manually defined. Using a methodology similar to Gao *et al*.^40^, we extracted 30 time-series related to the articulation process for each utterance. Among these, eight parameters are fixed values for a potential subsequent articulatory-to-audio conversion, and one is a binary estimation of speech vs. silence (‘subglottal pressure’). The remaining 21 parameters represent the trajectories of different articulatory muscles during the utterance production. However, one (‘vocal fold lower displacement’) was exactly the same as another (‘vocal fold upper displacement’) and thus discarded. The 20 remaining articulatory trajectories (in Supplementary Material) and the ‘subglottal pressure’ time-series were decimated to match the neural signal’s sampling rate. At this stage, the dataset was at the word level, meaning that we had a set (approximately 300 samples) of trajectories and neural data for each uttered word. The pipeline failed to generate proper trajectories for 20 word trials in total across all of the participant’s data (Table S1), these trials were discarded from further analysis. The sets of trajectories and neural data were concatenated back into continuous signals (Fig. 1B).

We trained a linear regression in a 10-fold cross-validation to predict each articulator separately for each channel and timeframe. Performance was calculated using the Pearson correlation between the original estimated and predicted articulatory trajectories over time. The folds were averaged to obtain the final score per channel, timeframe and articulator.

### Acoustic

The neural signal was further processed in the same way as for the articulatory representation, in 50 ms windows with a 10 ms frameshift. The audio signal was downsampled to 16 kHz and we extracted the speech spectrogram using the Short-Time Fourier Transform. This was also done in 50 ms windows with a 10 ms frameshift to ensure an alignment between the audio and neural features. The resulting spectrogram was compressed into a log-mel representation with 23 triangular filter banks (Fig. 1C). Thereafter, a linear regression was trained in a 10-fold cross-validation to predict each mel-bin (N=23) separately for each single channel and timeframe. Performance was calculated with the Pearson correlation between the original and the predicted mel-bin over time. The folds were averaged to calculate the final score for each channel, timeframe and mel-bin.

### Semantic

The neural signal was averaged across a larger window than for the other two representations, since semantic information is expected to have slower dynamics. This data was averaged in 200 ms windows with a 25 ms frameshift. The whole speech trial (2 seconds) was assigned to the cued word. Semantic embeddings (160 dimensional) were extracted for each Dutch word using a pre-trained *word2vec* model^41^. The embeddings were available for 85 of the 100 words, thus the 15 word trials without embeddings were discarded for each participant. As opposed to the previous two representations, we chose an encoding model for the semantic representation. A linear regression was trained to predict the neural timeseries for each recorded channel individually from the embeddings, in a 10-fold cross-validation (Fig. 1D). Performance was the Pearson correlation between the original and the predicted timeseries per window across word trials for each single channel. This way, variations in the average response to spoken words are evaluated as opposed to measuring how well the average response is captured. This procedure closely follows previous studies on semantic representations^2,38^.

### Statistical analysis

#### Significant channels

Randomized baseline performance was calculated for each of the representations individually. We applied a random circular shift procedure for the articulatory and acoustic representations. This procedure consisted of generating a random timepoint, between 10% and 90% of the data, on which the neural features were swapped. Then, the swapped data was correlated with the held-out ‘subglottal pressure’ time-series or each of the mel-bins and repeated 1000 times to estimate a distribution of chance correlation coefficients for every channel within every participant. The correlations across mel-bins were averaged before saving. Random permutations of the embedding vectors and re-training of the model were used to estimate a distribution of chance correlations for the semantic representation, this was also repeated 1000 times.

The significance threshold (*α* = 0.05) was set at the largest chance correlation across channels (articulatory and acoustic) or across channels and time windows (semantic) within each participant. Thereby, we used the max-*t* correction to correct for multiple comparisons^65^. The ‘subglottal pressure’ time-series was chosen in the articulatory pipeline as it distinguished overall speech from silence and yielded the strictest threshold. Any channel that had a correlation above the articulatory threshold in any of the articulators and timeframes was deemed significant for the articulatory representation. Thus, the results from individual articulators were pooled together to form one articulatory representation. The channel was significant if the average correlation across mel-bins for that channel was above the acoustic threshold in any of the timeframes for the acoustic representation. For the semantic representation, a channel was deemed significant if any of the time windows were above the semantic threshold.

#### Distribution metric

We took the coordinates (x, y, z) in MNI space from all significant channels within each hemisphere to calculate the geometric centroid for each representation. We then calculated the Euclidean distance between each channels’ coordinates and the centroid within each representation. Independent samples t-tests (Bonferroni-corrected for nine repeated tests) were used to calculate the statistical difference in distribution (i.e., distances) between the three representations within the hemispheres and between the left and right hemisphere within representations.

### Regions-of-interest

To find out how the speech representations are distributed within and across language areas, we did a region-of-interest (ROI) analysis in six language-related cortical areas (motor cortex, auditory cortex, superior temporal cortex, inferior frontal cortex, supramarginal cortex and the insula) and one subcortical area (hippocampus). The cortical areas included both gyri and sulci, a full description of which anatomical labels from the Destrieux/Fischl atlas we chose to include in each ROI can be found in the supplementary material (Table S2). Individual channels were selected based on the anatomical labels in individual space. We calculated the percentage of significant channels for each representation and their overlap out of the total number of channels present in that region. Other relevant regions mentioned in the introduction (the fusiform cortex, supplementary motor area, postcentral cortex and other subcortical areas) were not sufficiently sampled to be included as ROIs.

### Temporal dynamics

To leverage the temporal resolution of our recordings, we analyzed the temporal dynamics of the results. The number of significant channels and their correlations were evaluated within each time-point or time-frame. Since the semantic analysis is trial-based, the time-points go from 0 ms (word cue) to 1875 ms (end of shortest trial). Since the motoric and acoustic analysis is based on the entire recording, the time-frames represent the misalignment between the neural and target data from -200 ms to +200 ms. The same cross-validation and statistical procedures were used as in the main analysis.

### Acoustic contamination

Electrophysiological recordings can potentially be contaminated by the acoustics of overt speech, which can be detected with a significant correlation in the spectral energy between the recorded neural and audio data^66^. None of the 15 participants had a significant correlation (all *p* > 0.01), indicating a chance of less than 1% of acoustic contamination.

## Data availability

The preprocessed data is publicly available at https://doi.org/10.17605/OSF.IO/QZWSV and code can be found on https://github.com/neuralinterfacinglab/SpeechRepresentations.

## Acknowledgements

This publication is part of the project INTENSE (with project number 17619 of the research programme NWO Crossover Programme) which is (partly) financed by the Dutch Research Council (NWO). C.H. acknowledges funding by the Kavli Foundation.

## Author contributions statement

C.H. and P.K. designed the experiment, C.H, M.C.O. and M.V. collected the data, J.V. and Y.G. generated the articulatory trajectories, M.V. and C.H. ran all other analyses and wrote the manuscript. All authors reviewed the manuscript and declare no competing interests.

## Supplementary Material

### Included articulatory trajectories

Subglottal pressure, Vocal fold upper displacement, Chink area, Relative amplitude, Horizontal hyoid position, Vertical hyoid position, Jaw position, Jaw angle, Lip protrusion, Vertical lip distance, Velum shape, Velum opening, Horizontal tongue body center position, Vertical tongue body center position, Horizontal tongue tip position, Vertical tongue tip position, Horizontal tongue blade position, Vertical tongue blade position, Tongue side elevation (posterior), Tongue side elevation (middle), Tongue side elevation (anterior).

**Table S1.**
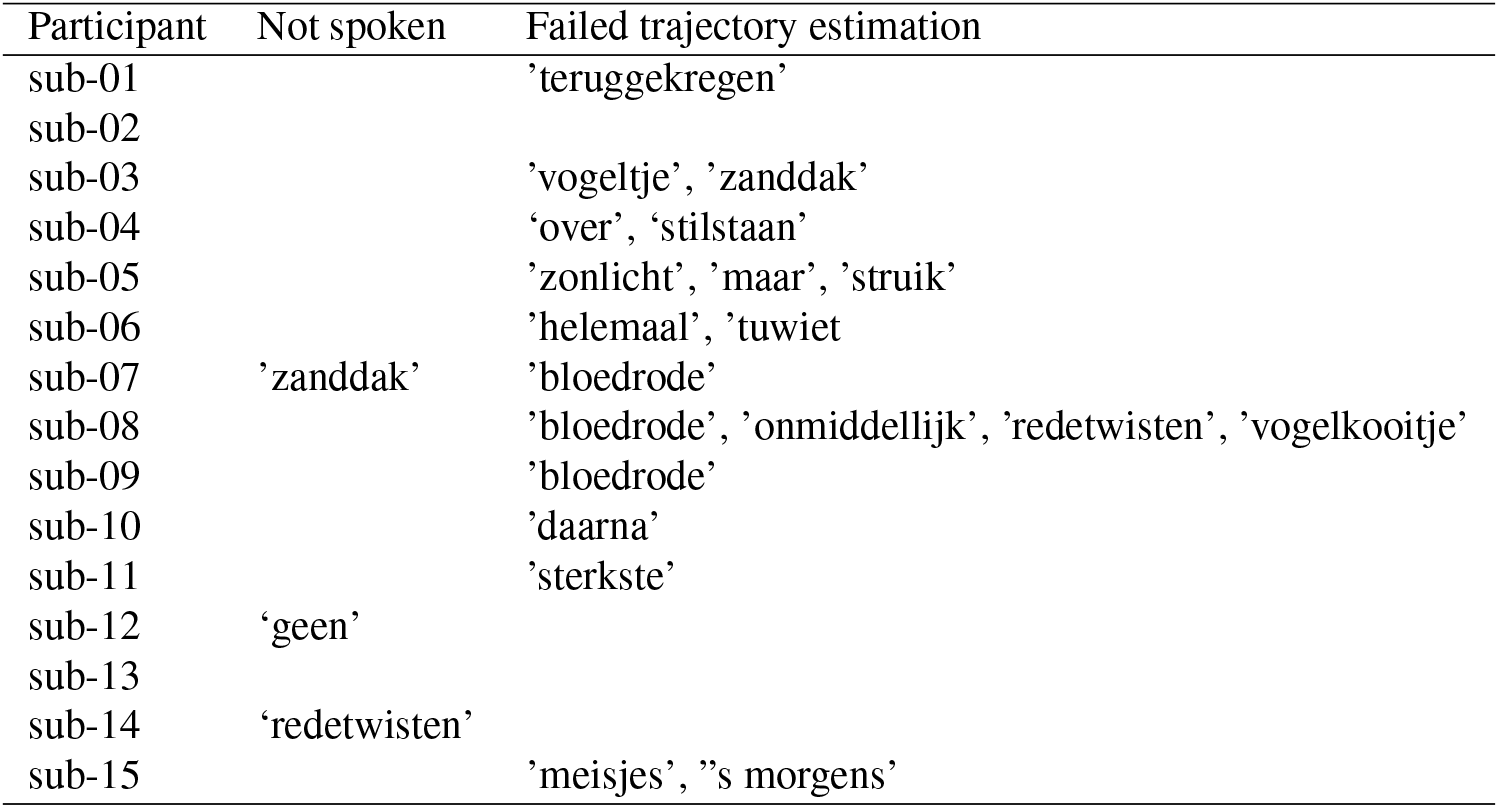
Discarded articulatory word trials for each participant and the reason for exclusion (not spoken or failed trajectory estimation).

**Table S2.** Included Destrieux/Fischl atlas labels per region-of-interest. The same labels were applied to both hemispheres.

| Region-of-interest | Labels |
| --- | --- |
| Auditory cortex | 'S_temporal_transverse', 'G_temp_sup-G_T_transv', 'G_temp_sup-Plan_tempo' |
| Inferior frontal cortex | 'G_front_inf-Opercular', 'G_front_inf-Orbital', 'G_front_inf-Triangul', 'G_orbital', 'S_orbital-H_Shaped', 'S_suborbital', 'S_front_inf', 'S_orbital_lateral', 'Lat_Fis-ant-Vertical', 'Lat_Fis-ant-Horizont' |
| Motor cortex | 'S_central', 'G_and_S_subcentral', 'G_precentral', 'S_precentral-inf-part' |
| Superior temporal cortex | 'G_temp_sup-Lateral', 'S_temporal_sup', 'G_temp_sup-Plan_polar' |
| Supramarginal cortex | 'G_pariet_inf-Supramar', 'Lat_Fis-post' |
| Insula | 'S_circular_insula_inf', 'S_circular_insula_sup', 'G_insular_short', 'G_Ins_lg_and_S_cent_ins' |
| Hippocampus | 'Hipp' |

**Figure S1.**
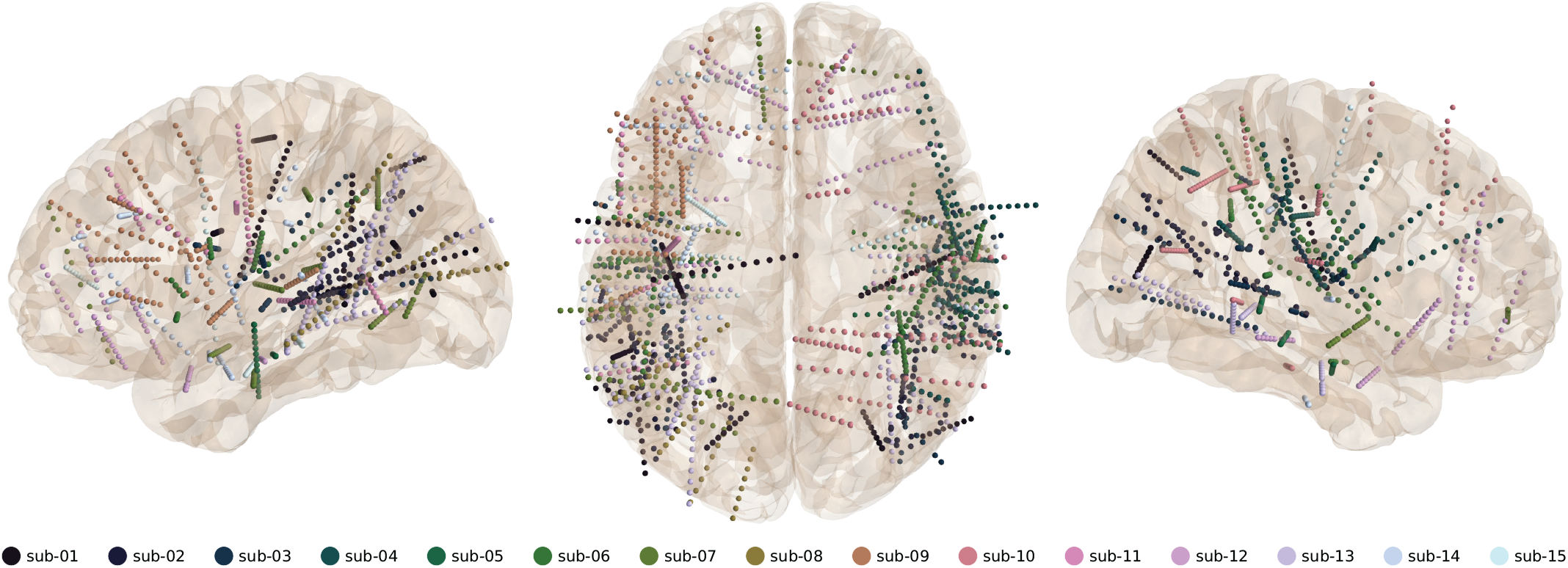
Electrode locations. Colors represent the participants.

**Figure S2.**
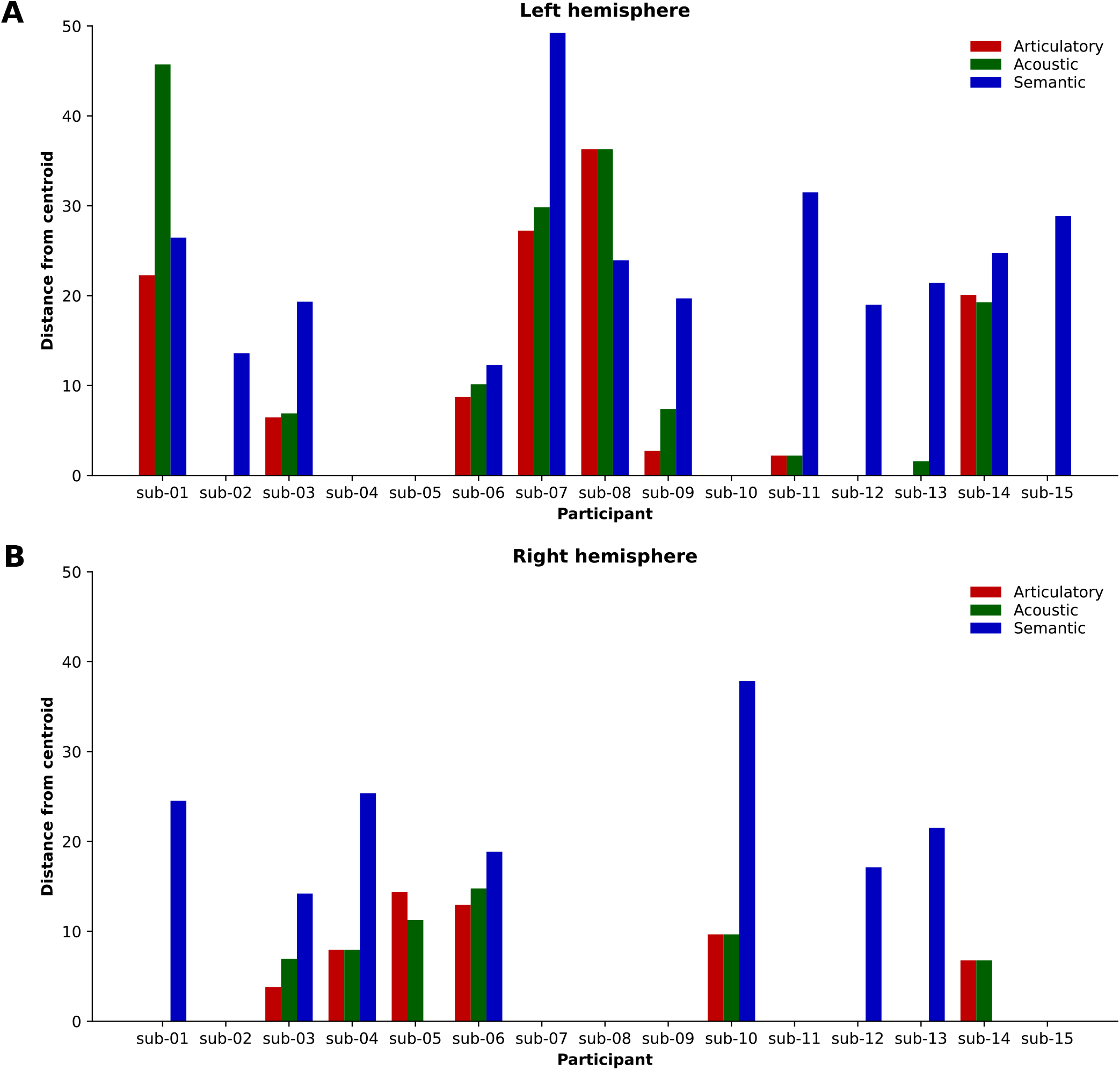
Distribution between representations within individuals. The distribution in calculated using the mean Euclidean distance between each significant channel and their centroid for each representation within individuals. **A)** Distances within the left hemisphere. **B)** Distances within the right hemisphere.

**Figure S3.**
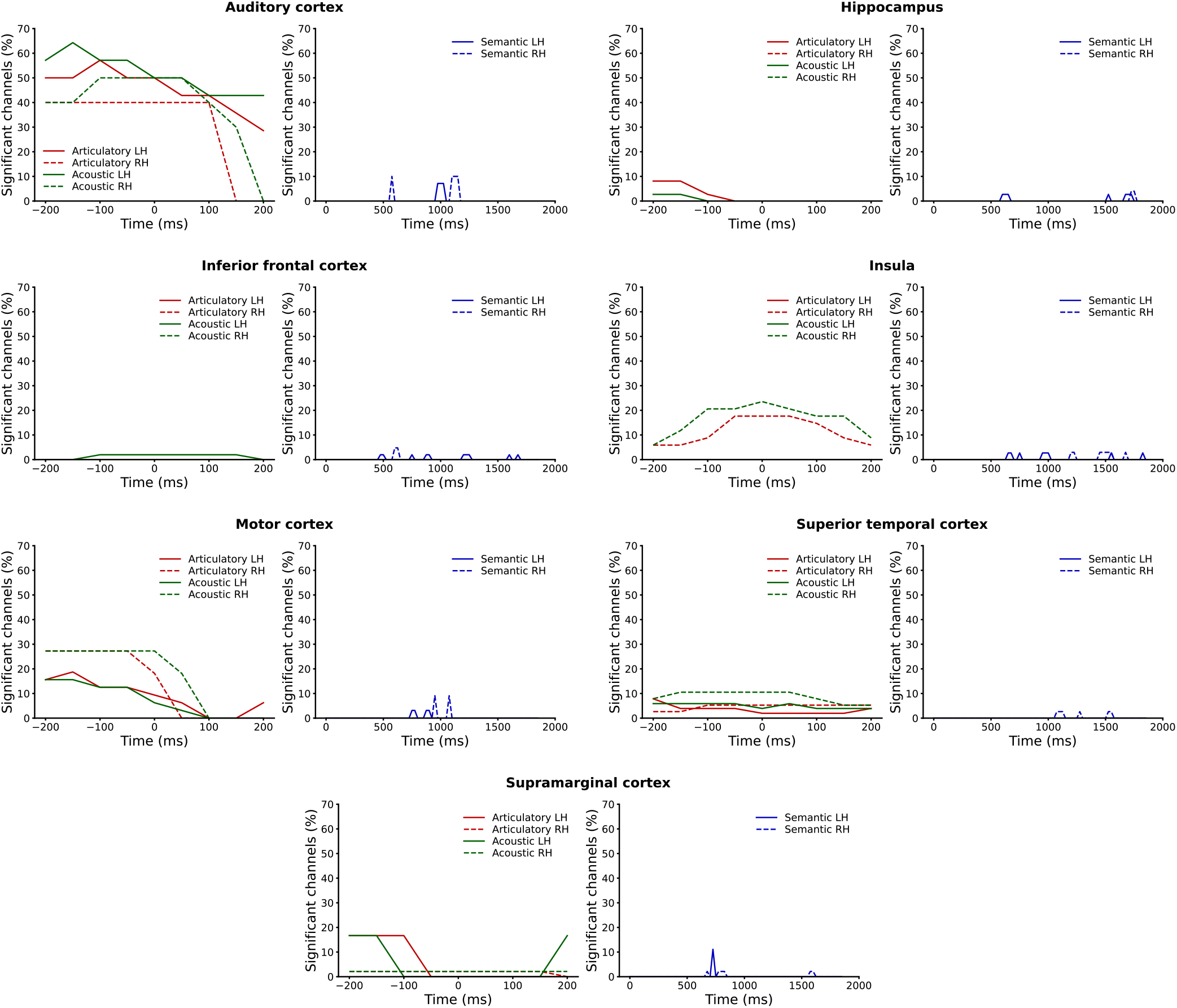
Temporal dynamics for all seven regions-of-interest. The percentage of significant channels within each region for each timeframe or timepoint is plotted to facilitate comparisons.

